# Microglial elimination of dopamine D1 receptors defines sex-specific changes in nucleus accumbens development and social play behavior during adolescence

**DOI:** 10.1101/211029

**Authors:** Ashley M. Kopec, Caroline J. Smith, Nathan R. Ayre, Sean C. Sweat, Staci D. Bilbo

## Abstract

Adolescence is a developmental period in which the mesolimbic dopaminergic ‘reward’ circuitry of the brain, including the nucleus accumbens (NAc), undergoes significant developmental plasticity and neural circuit maturation. Dopamine D1 receptors (D1rs) in the NAc have recently been demonstrated to be critical modulators of social behavior, but how these receptors are regulated in adolescence to mediate social behavior is not well understood. In this report, we used multi-plexed immunohistochemistry with volumetric reconstructions, co-immunoprecipitation, *ex vivo*, and *in vivo* stereotaxic, microglial manipulation, and social behavior assessment to demonstrate that microglia and complement-mediated phagocytic activity shapes sex-specific NAc development. Moreover, we report for the first time that microglia-mediated phagocytosis is required for natural developmental changes in behavior, specifically, adolescent male social play behavior. These data have broad implications for understanding how experience interacts with the developing reward circuity, sex-specific responses to stimuli in adolescence, and how neuropsychiatric disorders may arise in a sexually dimorphic manner.

## Introduction

Adolescence is a developmental stage characterized by increased exploration, risk taking, and social interaction.^1,2^ During this period, the mesolimbic dopaminergic ‘reward’ circuitry of the brain, which includes the ventral tegmental area (VTA), prefrontal cortex (PFC), and nucleus accumbens (NAc), undergoes significant developmental plasticity and neural circuit maturation.^3^ Behaviors which engage the reward circuitry during this window of active developmental neuroplasticity can disproportionately impact brain development and, consequently, later life behavior.^4^

A behavioral hallmark of adolescence is increased peer-centered social interaction.^2,5^ Neuropeptides commonly associated with social behavior, like oxytocin, endocannabinoids, and opioids, have been demonstrated to modulate social behavior when manipulated locally in the amygdala, NAc, and VTA, in rodents,^6–11^ as well as intra-nasally in humans.^12^ Interestingly, access to social experience can be used in lieu of addictive drugs to induce conditioned place preference, operant conditioning, or maze performance,^13,14^ and social interactions can modulate addiction-like behaviors, acting to enhance or diminish drug-seeking behaviors in difference contexts.^15^ Futhermore, NAc dopamine transients are induced by brief social interactions in both adult and adolescent rats; however, adult dopamine responses habituate, while adolescent dopamine responses persists in subsequent peer interactions.^16^ Taken together, these data suggest that social interaction itself is a highly rewarding/motivating experience that engages the dopaminergic reward circuitry.

Dopamine receptors in the NAc have recently been demonstrated to be required for social play behavior in adolescent rats,^17^ and two seminal studies have used optical and genetic tools to define a hypothalamic-VTA-NAc circuit underlying social behavior. Oxytocin innervation from the paraventricular nucleus of the hypothalamus (PVN) to the VTA increases VTA neuron excitability, and initiates dopamine release into the NAc, promoting pro-social behavior.^10^ Remarkably, the VTA-NAc dopaminergic response was mediated specifically by NAc dopamine D1 receptors (D1rs), and not dopamine D2 receptors (D2rs) in adult female mice.^18^ Moreover, NAc D1r, but not D2r signaling both basally and during social interaction was capable of predicting individual susceptibility to depression induced by a social defeat paradigm.^19^ Interestingly, several subtypes of dopamine receptors, including D1rs, are downregulated (via unknown mechanisms) during adolescence in the PFC and dorsal striatum.^20–24^ However, there are conflicting reports of D1r regulation during adolescent development in the NAc: some reports conclude dopamine receptors are not downregulated in the NAc in adolescence,^20^ while other reports conclude that they are, indeed, downregulated.^21,23^ Thus, the data collectively suggest that NAc D1rs are a critical mediator of social behavior, and potentially also the interpretation of social interactions, both of which are critical features of adolescent development. Two complementary open questions remain: (i) how does the dopamine system in the NAc develop? and (ii) does this development modulate social behavior in adolescence?

Microglia, the resident immune cells of the brain, are critical mediators of neural circuit development in certain brain regions, e.g. in synaptic pruning and refinement.^25^ For instance, in the developing retinogeniculate nucleus of male mice, the classical immune complement system, including complement protein C3, ‘tags’ synapses for elimination^26^. Microglia, which exclusively express C3 receptor (C3R, often referred to as CD11b), recognize this tag, phagocytose, and lysosomally degrade the synapse, which results in the developmental refinement of ocular inputs.^26,27^ We thus sought to determine if microglia and immune signaling participate in dopaminergic development via D1r elimination in the NAc during adolescence. Furthermore, we wished to determine whether microglial-mediated circuit refinement can account for normal developmental changes in behavior, which as of yet has not been established.

Herein, we report sex-specific microglia and complement-mediated phagocytosis and elimination of dopamine D1 receptors in the adolescent NAc. While D1rs are downregulated in both the male and female NAc at different ages, microglia and complement C3-mediated phagocytic elimination is only engaged in males. Moreover, we report for the first time that microglia-mediated phagocytosis is required for natural developmental changes in behavior, specifically, adolescent social play behavior in males. These data have broad implications for understanding how experience interacts with the developing reward circuity, sex-specific responses to stimuli in adolescence, and how neuropsychiatric disorders may arise in a sexually dimorphic manner.

## Results

### Dopamine D1 receptor downregulation over adolescent development is associated with complement C3 and microglia in the male, but not the female, nucleus accumbens

Our first goal was to determine the pattern of D1r expression over the course of male and female adolescent development in rats. We assessed tissue at four ages: postnatal day 20 (<u>P</u>20) representing peri-adolescent animals, P30 representing early adolescence, P38 representing mid-adolescence, and P54 representing late adolescence. We used D1r immunoreactivity to define our regions of interest, and focused on the dorso-medial area of the anterior NAc (Supp. Fig. 1A). Immunofluorescent analysis revealed significant D1r downregulation in male rats between early-mid adolescence and (Fig. 1A; Table 1A) and in female rats from peri-early adolescence (Fig. 1E; Table 1E), which was confirmed in a separate group of animals using chromogenic immunohistochemistry (Supp. Fig. 1B,C).

**Table 1.**
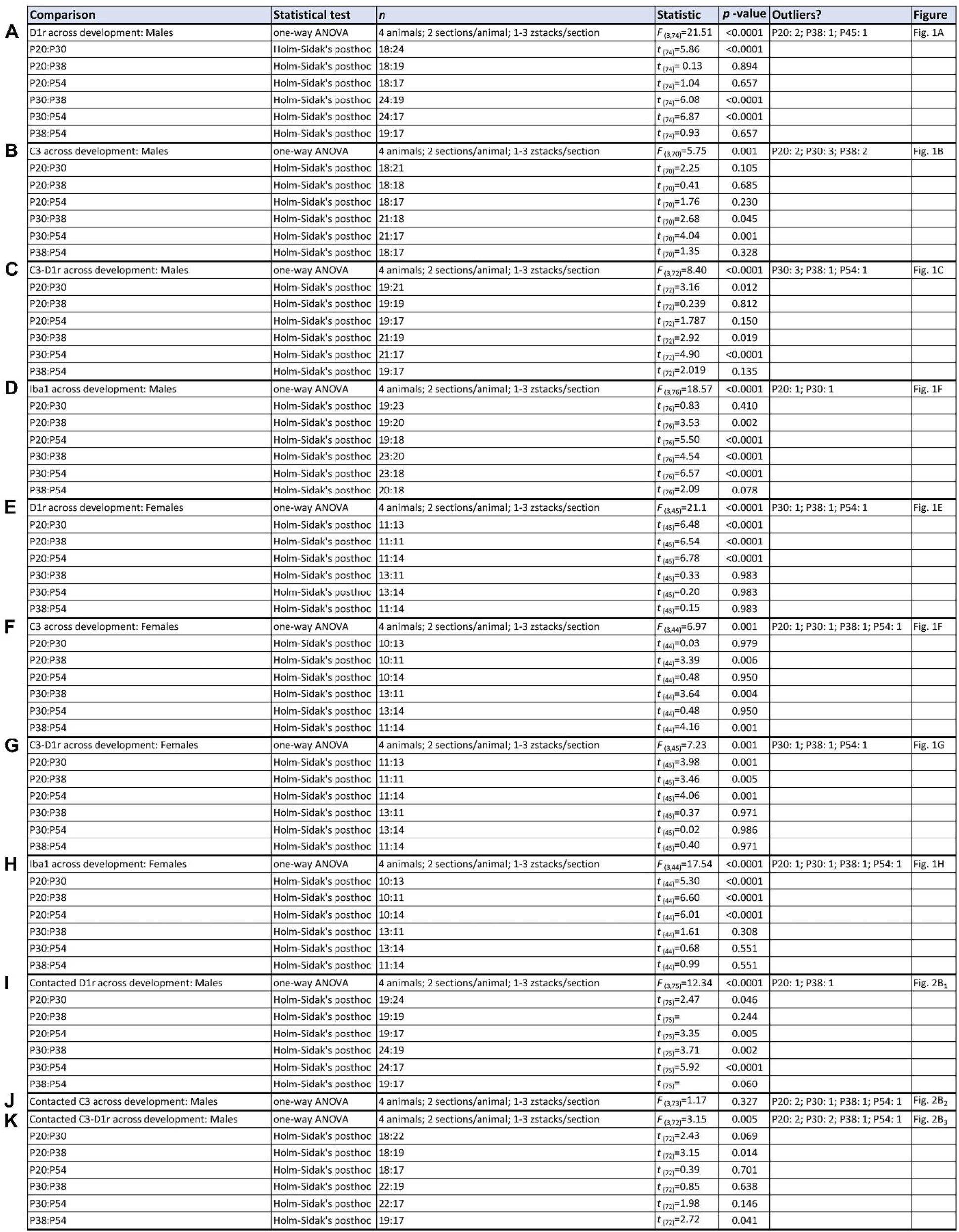
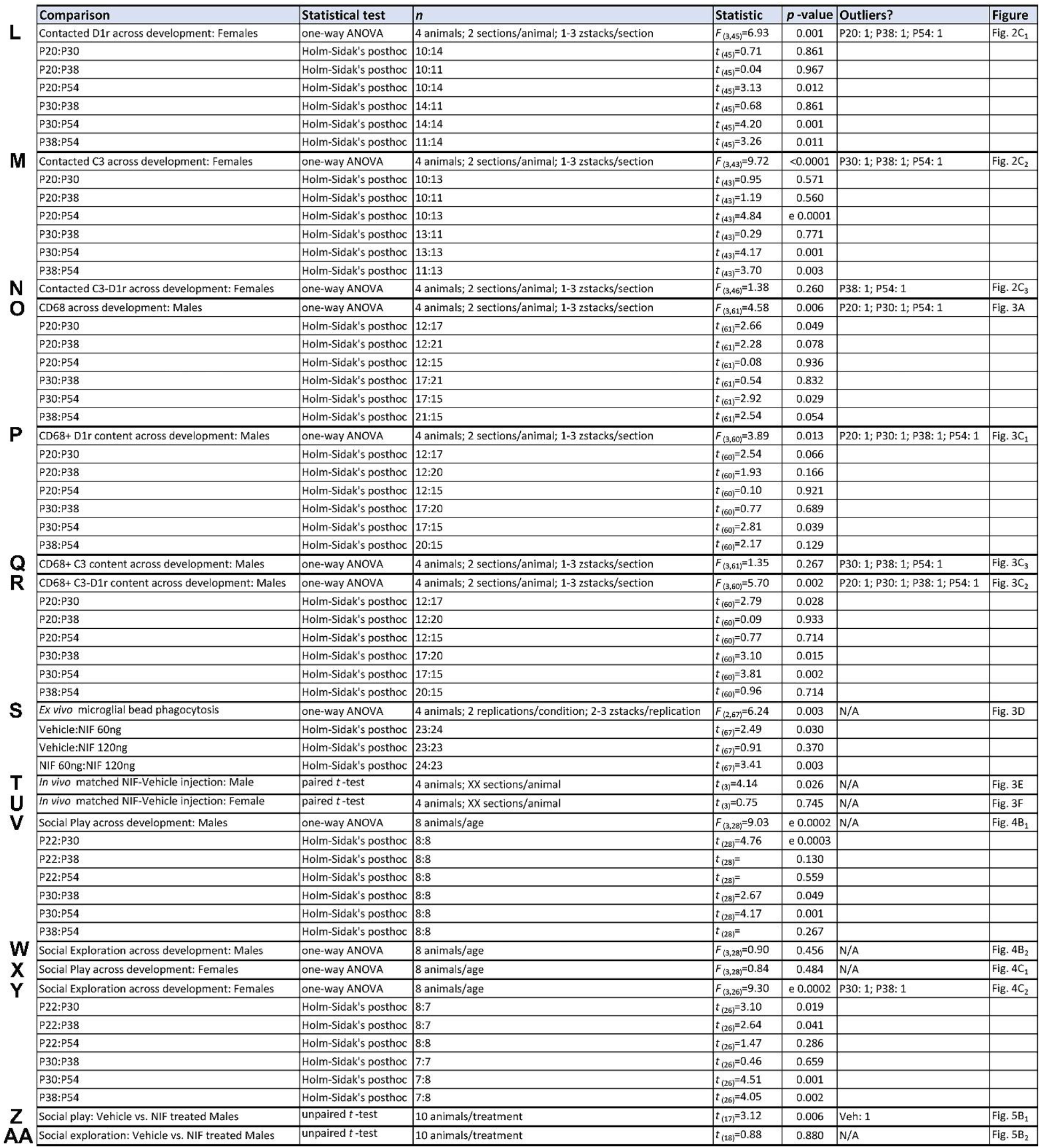
Detailed Statistics. Statistical details for every analysis.

**Fig. 1.**
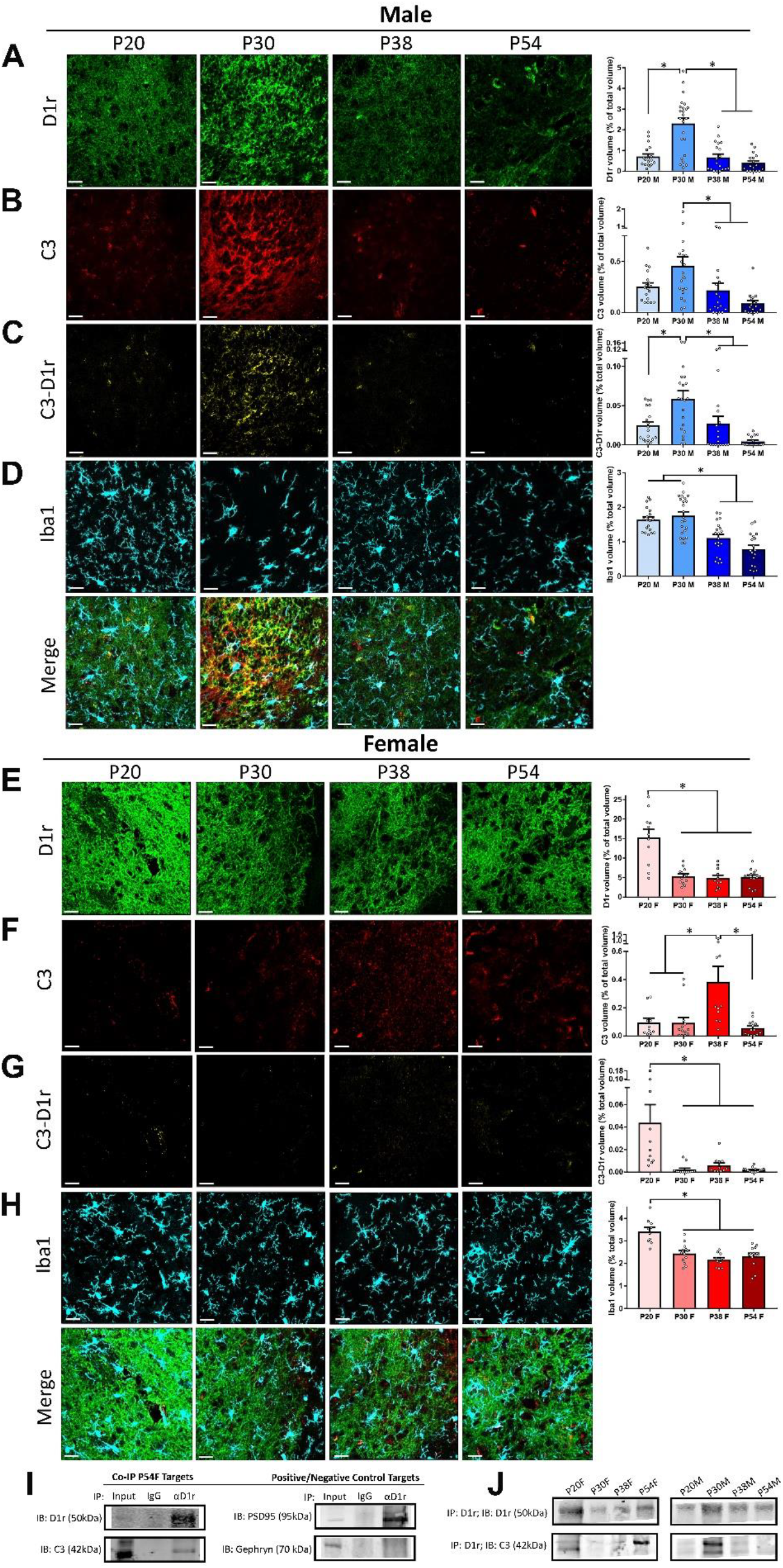
D1r downregulation in the NAc occurs at sex-specific ages and is associated with the phagocytosis ‘tag’, complement protein C3, in males, but not females. **(A-H)** Triple-label immunohistochemistry for D1r, C3, and Iba1 was performed and protein expression levels for each target and for a C3-D1r colocalization channel were normalized for z-stack size and analyzed with one-way ANOVAs and Holm-Sidak’s post-hoc comparisons. Representative images precede histograms and merged images are below; scale bars are equal to 20µm. In both males and females, *n*=4 animals/age/sex. Histograms portray the mean +/− SEM with individual data points overlaid. Significant (*p*<0.05) comparisons are delineated with an asterisk. All statistics are located in Table 1. <u>Males</u>: **(A)** D1r levels transiently increased at P30 (Table 1A); **(B)** C3 levels decreased after P30 (Table 1B); **(C)** Levels of C3 colocalized with D1r transiently increased at P30 (Table 1C). **(D)** Iba1 levels decreased after P30 (Table 1D). <u>Females</u>: **(E)** D1r levels decreased after P20 (Table 1E); **(F)** C3 levels transiently increased at P38 (Table 1F); **(G)** C3-D1r levels decreased after P20 (Table 1G); **(H)** Iba1 levels decreased after P20 (Table 1H). **(I-J)** To determine if the C3 cleavage product iC3b (~42kDa), thought to act as an opsonin, could physically associate with D1rs, αD1r co-immunoprecipitation (Co-IP) was performed on NAc samples. **(I)** Assessment of the P54 female sample detected D1r in pre-IP input (albeit at low levels) and enrichment for D1r after αD1r, but not αIgG control IP. Similarly, iC3b and PSD95 were detected in Input and αD1r IP eluates, while gephyrin was only detected in Input samples. **(J)** One sample/sex/age was subjected to αD1r Co-IP and eluates were assessed for D1r and iC3b. Co-IP confirmed that D1r and iC3b could, indeed, physically associate (most notably: in males, P30 and P38; in females P20 and P54).

We next determined if complement C3 and microglia are involved in D1r downregulation, consistent with their role in synaptic pruning and circuit refinement in other developing brain regions. In the peripheral immune system, C3 is proteolytically processed into an opsonin, which behaves as a ‘tag’ to target and augment phagocytic activity by C3 receptor (C3R, also referred to as CD11b)-expressing macrophages, a process referred to as opsonization.^28,29^ We assessed both total C3 levels as well as C3 colocalization with D1r. If C3 is serving as an opsonin recognizable by C3R/CD11b on the brain’s macrophages (i.e. microglia) to mediate D1r elimination, then C3-D1rs would be expected to be a preferential target of microglial phagocytic activity. Consistent with this notion, total C3 levels (Fig. 1B; Table 1B) and C3-D1r levels (Fig. 1C; Table 1C) are high at P30 in males, prior to D1r downregulation. However, in females, C3 levels were highest at P38 (Fig. 1F; Table 1F), while C3-D1r levels were highest at P20 (Fig. 1G; Table 1G). Interestingly, in both males and females, decreased Iba1 expression (a ubiquitous protein on microglia) was observed concurrent with D1r elimination (Fig. 1D,H; Table 1D,H). In the peripheral immune system, the C3-derived opsonin is iC3b, a cleavage product of C3.^29^ Because our immunofluoresent antibody was pan-C3 and will detect all forms (cleaved or uncleaved) of C3, we performed D1r co-immunoprecipitation on NAc samples to determine if iC3b (identified by its unique molecular weight ~42kDa)^30^ could be detected in complex with D1r. D1r has been reported to associate with the post-synaptic scaffolding molecule of excitatory synapses, PSD-95;^31,32^ conversely, D1r does not associate inhibitory synapses (specifically, GABA^A^ receptors) in hippocampus.^33^ Thus, we assessed immunoprecipitation of PSD-95 and gephyrin, an inhibitory synapse post-synaptic scaffold, as putative positive and negative controls in addition to D1r and C3. The anti-D1r antibody used for immunofluorescent analysis was capable of precipitating C3 and PSD95, but not gephyrin, while the Input sample (prior to Co-IP procedures) was positive for all proteins and control IgG had no association (Fig. 1I). Indeed, at several different ages in both males and females there is evidence for iC3b association with D1r (Fig. 1J). There is another intriguing hypothesis derived from these data, though it should be interpreted conservatively. The data replicate peak levels of D1r at P30 and P20 in males and females, respectively, and while iC3b was detectable in all eluates, the highest level of immunoprecipitated iC3b in males corresponded with highest D1r levels at P30, prior to D1r downregulation. Contrarily, the highest level of immunoprecepitated iC3b in females was at P54, and thus not associated with the highest levels of D1r (Fig. 1J). Taken together, these data may suggest that while an iC3b-D1r tagging mechanism resulting in D1r elimination may be employed in male adolescent NAc development, iC3b and D1r downregulation are uncoupled in female development.

If C3 (and/or its cleavage product) is being utilized as a tag for subsequent phagocytic activity in males specifically, then microglia, which exclusively express C3R/CD11b, should contact C3-D1r complexes and mediate their elimination via phagocytosis in the male, but not female NAc. To assess this, we used volumetric reconstruction of Iba1 immunoreactivity to measure the volume of D1r, C3, and C3-D1r in contact with, or inside, microglia (Fig. 2A). Because all protein expression assessed changed significantly changed (D1r, C3, C3-D1r, and Iba1; Fig. 1), increased microglial contact could result from three mechanisms: (i) increased total D1r (or C3/C3-D1r) levels increase the likelihood of microglial contact by virtue of the increased space they occupy; (ii) increased Iba1 levels may increase levels of D1r contact by the same principle; or (iii) microglia are actively contacting D1rs irrespective of fluctuating D1r or Iba1 levels. Our hypothesis presupposes that microglia are actively contacting D1rs due to recognition of the D1r-associated C3 tag; thus to test this hypothesis specifically we normalized the volume of contacted protein for both (a) total protein of the target and (b) Iba1 levels (see *Methods*). In males, both D1rs (Fig. 2B_1_; Table 1I) and C3-D1rs (Fig. 2B_3_; Table 1K) are, indeed, maximally contacted by microglia prior to D1r downregulation. Male microglial contact of C3 does not change over time (Fig. 2B_2_; Table 1J), potentially indicative of constitutive microglial C3R – C3 interactions. However, in females, microglia maximally contact D1rs (Fig. 2C_1_; Table 1L) and C3 (Fig. 2C_2_; Table 1M) at P54, rather than prior to D1r elimination at P20, which is consistent with a greater association of C3 with D1r at P54 suggested by the immunoprecipitation data. Interestingly, there is no significant difference in C3-D1r contact (Fig. 2C_3_; Table 1N), indicating that while there may be meaningful C3-microglia, C3-D1r, and/or D1r-microglia interactions in the female NAc, unlike the males, they are not associated with D1r regulation at the ages assessed. Taken together, the data suggest that in males, complement C3 (and perhaps more specifically iC3b) associates with D1r and results in microglial recruitment at P30 and subsequent elimination of D1rs at P38; in females, D1r downregulation does not appear to be contingent on C3 or microglia.

**Fig. 2.**
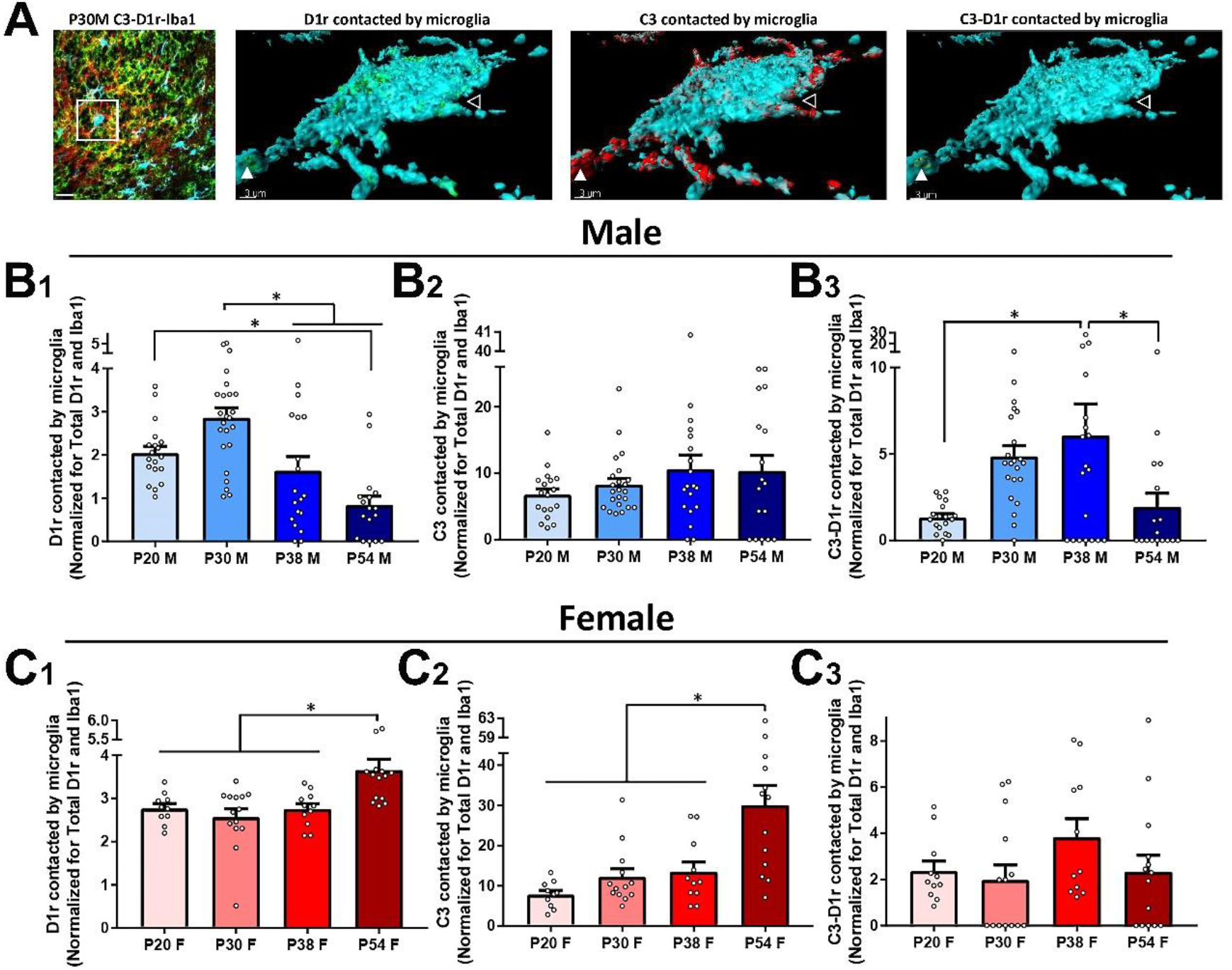
Microglia contact NAc D1r/C3-tagged D1rs prior to overt D1r downregulation in males, but not females. Triple-label immunohistochemistry in Fig. 1 were analyzed using a volumetric reconstruction of Iba1 (i.e. microglia) as a mask to determine the volume of D1r, C3, and C3-D1r physically associated with (either in contact with or inside) microglia. Masked volumes were normalized for total protein of the target and total Iba1, to control for ‘passive’ interactions by virtue of increased microglia or increased target protein. Data are analyzed with one-way ANOVAs and Holm-Sidak’s post-hoc comparisons. In both males and females, *n*=4 animals/age/sex. Histograms portray the mean +/− SEM with individual data points appearing overlaid. Significant (*p*<0.05) comparisons are delineated with an asterisk. All statistics are located in Table 1. **(A)** Representative 2D triple-label immunohistochemistry (scale bar 20µm), with enlarged 3D representation (scale bar 3µm) of the selected area demonstrating (left-right) D1r, C3, and C3-D1r contacted by/internal to the microglia. Open arrow head indicates a region of the microglia where both D1r and C3 are located, but not co-localized, demonstrating the sensitivity of co-localization channels. Closed arrow head indicates a region of the microglia where D1r and C3 co-localize (C3-D1r). <u>Males</u>: **(B_1_)** D1rs are maximally contacted by microglia (irrespective of changing D1r and Iba1 levels) at P30, prior to their elimination (Table 1I). **(B_2_)** C3 contact does not change over time (Table 1J), and **(B_3_)** C3-D1r contact is maximal between P30-P38 (Table 1K). Females: **(C_1_)** D1rs are maximally contacted by microglia at P54, not at P20, prior to their elimination (Table 1L). **(C_2_)** C3 contact is also increased at P54 (Table 1M), while **(C_3_)** C3-D1r contact does not change over development (Table 1N).

### Adolescent regulation of D1rs is mediated by microglia- and complement C3-mediated phagocytosis in the male, but not female NAc

Our data thus far support the notion that the complement system is mediating D1r elimination in the developing NAc in males, but not females. CD68 is a lysosomal protein in microglia and thus has been used as a marker to examine increased phagocytic activity, such as synaptic pruning.^26,27^ Thus, we first measured the expression levels and content of CD68+ microglial lysosomes over the course of adolescent development in males. CD68 expression is increased at P30-38 (Fig. 3A; Table 1O), suggesting increased phagocytic activity in microglia. Association with CD68+ lysosomes is an inherently active process achieved via engulfment (Fig. 3B), and thus we next measured the content of CD68+ lysosomes by calculating the percentage of D1r, C3, and C3-D1r contained within CD68+ lysosomes as a percentage of total D1r, C3, and C3-D1r protein. The percentage of D1r (Fig. 3C_1_; Table 1P) and C3-D1r (Fig. 3C_3_; Table 1R) content within lysosomes was highest between P30 and P38. In accordance with no change in microglial contact of C3 over development, there was also no change in C3 levels associated with CD68+ lysosomes (Fig. 3C_3_; Table 1Q).

**Fig. 3.**
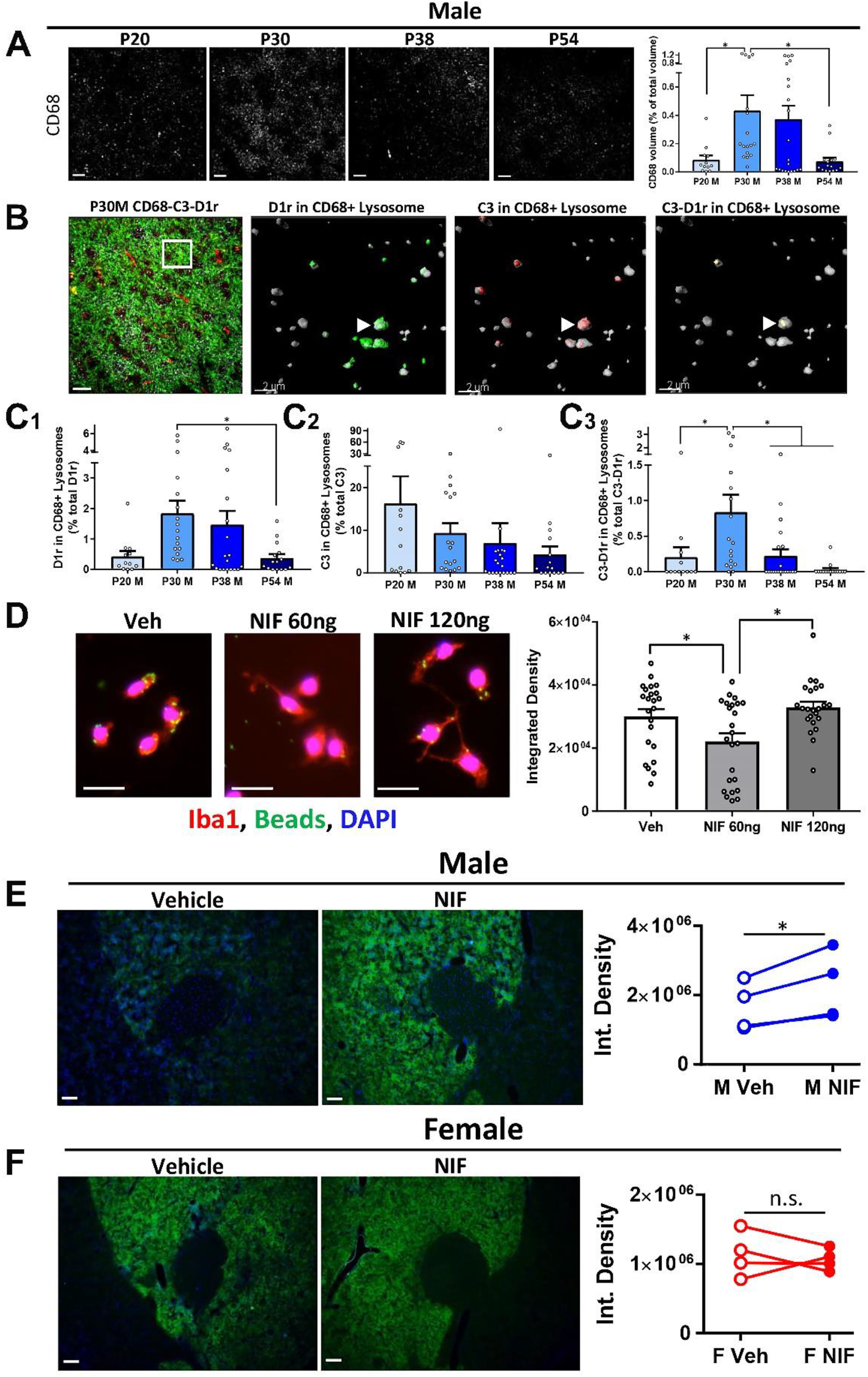
Microglia phagocytose and lysosomally eliminate C3-tagged D1rs during adolescent NAc development in males. **(A-C)** Triple-label immunohistochemistry for CD68, D1r, and C3 was performed on male NAc (*n*=4 animals/age, with 12-20 observations in each group) to assess CD68 levels over development and CD68+ lysosomal content of D1r, C3, and C3-D1r. Because association with a lysosome is an inherently active process (i.e. engulfment by microglia and subsequent shuttling to lysosome), lysosomal content was normalized only for total protein of the target. Data were analyzed with one-way ANOVAs and Holm-Sidak’s post-hoc comparisons. Histograms portray the mean +/− SEM with individual data points overlaid. Significant (*p*<0.05) comparisons are delineated with an asterisk. All statistics are located in Table 1. **(A)** CD68 levels, which correlate with phagocytic activity in microglia, transiently increase at P30. Representative images precede the histogram (Table 1O; scale bar 20µm). **(B)** Representative 2D triple-label immunohistochemistry (scale bar 20µm), with enlarged 3D representation (scale bar 2µm) of the selected area demonstrating (left-right) D1r, C3, and C3-D1r content in CD68+ microglial lysosomes. Closed arrow head indicates a CD68+ lysosome where D1r and C3 co-localize (C3-D1r).**(C_1_)** D1r content in CD68+ lysosomes is highest at P30-38, irrespective of overt changes in D1r levels (Table 1P). **(C_2_)** C3 content does not change over development (Table 1Q), and **(C_3_)** C3-D1r content is transiently elevated at P30 (Table 1R). **(D-F)** Neutrophil inhibitor factor (NIF), a peptide that binds specifically to the CD11b subunit of C3 receptor, was assessed for its efficacy in reducing microglial phagocytic activity *ex vivo* and *in vivo*. For *ex vivo* experiments, *n*=4 (all adult female) and 2 replications/experimental condition. For *in vivo* experiments, n=4 animals/sex and data analyzed are the average value of 4-7 observations in males (per hemisphere) and 5-6 observations in females. Significant (*p*<0.05) comparisons are delineated with an asterisk. All statistics are located in Table 1. **(D)** 60ng, but not 120ng NIF inhibited microglial phagocytosis of fluorescent beads (Table 1S). Data are analyzed with one-way ANOVAs and Holm-Sidak’s post-hoc comparisons. Histograms portray the mean +/− SEM with individual data points appearing overlaid. Representative images precede histograms. **(E-F)** NIF was microinjection into one hemisphere and vehicle microinjection into the contralateral hemisphere in P30 males or P22 females (both ages prior to D1r elimination), and then within-animal D1r levels were compared at P38 and P30 in males and females, respectively. **(E)** NIF treatment increased D1r levels in males (Table 1T), but not **(F)** females (Table 1U). Data are analyzed with paired 2-tailed *t*-tests. Histograms portray the within-animal change in D1r levels in vehicle and NIF-treated hemispheres. Representative images precede histograms.

In adolescent males, the data thus far converge to suggest D1r downregulation in the NAc results from C3 tagging, microglial recognition, phagocytosis, and lysosomal degradation. If so, blocking the ability of microglial C3R to recognize and bind the C3 tag associated with D1r should disrupt their phagocytic elimination of D1rs, thus augmenting D1r expression. Neutrophil inhibitor factor (NIF) is a well-characterized canine hookworm peptide that binds specifically to the CD11b subunit of C3R, occluding its ability to bind its natural ligands, including iC3b.^34–36^ NIF has been used systemically to (i) improve stroke outcomes in a rat model,^37^ (ii) treat diabetic retinopathy in mice,^38^ and (iii) as an exploratory treatment for stroke in humans.^39,40^ However, to our knowledge, whether NIF can inhibit phagocytosis by microglia or function locally in the brain has not been assessed. We first tested whether NIF can inhibit phagocytosis in microglia *ex vivo*. Microglia were isolated from whole, adult female brains and treated with media containing either vehicle, 60ng NIF (reconstituted according to manufacturer’s recommendations at 200µg/mL; 1x), or 120ng NIF (reconstituted at 400µg/mL; 2x) on glass coverslips. Fluorescent carboxylate microspheres (hereafter referred to as “beads”) were applied to induce phagocytic activity. Phagocytic activity was significantly impaired when microglia were incubated with 60ng, but not 120ng of NIF (Fig. 3D; Table 1S). Thus, to block microglial C3R – C3 association and consequent phagocytosis *in vivo*, we microinjected 60ng NIF (in 300nL) into the dorsomedial NAc of one hemisphere, and vehicle into the contralateral hemisphere of P30 males (Supp. Fig. 2). At this age, microglial contact of C3-D1rs is high, but D1r elimination has not been fully accomplished, and if D1r elimination occurs by virtue of microglial C3 receptor – C3 interaction and microglial phagocytosis, D1r levels should be elevated in NIF-treated hemispheres relative to vehicle-treated hemispheres within the same animal at a later age (i.e. P38). Moreover, to confirm that female D1r elimination *is not* dependent on the same mechanism as males, we performed the same experiment in P22 females, prior to their D1r downregulation at P30. Indeed, in males, D1r levels in NIF-treated hemispheres were significantly elevated relative to vehicle-treated hemispheres (Fig. 3E; Table 1T), while there was no difference in D1r levels between NIF and vehicle-treated hemispheres in females (Fig. 3F; Table 1U). These data demonstrate for the first time a sex-specific immune mechanism regulating dopaminergic NAc development during adolescence.

### Social behavior exhibits sex-specific patterns in adolescence that require microglial C3R-C3 interactions in males, but not females

We next sought to determine if D1r downregulation, and thus the mechanisms which regulate this process, could have consequences for normal developmental changes in adolescent behavior. Social behavior, and in particular, social play behavior is known to peak during adolescence in males and requires dopamine signaling in the NAc.^17^ Moreover, recent dissection of the neural circuitry supporting pro-social behavior has implicated D1rs in the NAc as a key substrate.^10,18^ Thus, male and female rats were single-housed for 24 hours, a period of isolation known to increase social interaction and play without inducing long-lasting detrimental effects,^41–43^ and then a novel age- and sex-matched rat was introduced into the home cage and their interactions were recorded for 10 minutes (Fig. 4A). Both social play and social exploration (i.e. non-play social behavior consisting of allogrooming and sniffing of the conspecific) initiated by the experimental animal was scored by an experimenter blind to age and sex conditions. Strikingly, engagement in social play behavior mirrored the pattern of D1r expression in males: social play transiently peaked in early adolescence, and was down to peri-adolescent levels by late adolescence (Fig. 4B_1_; Table 1V), with no changes in social exploration (Fig. 4B_2_; Table 1W). In contrast, play levels did not significantly change over the ages assessed in females (Fig. 4C_1_; Table 1X). Rather, female social exploration increased in early adolescence and then decreased again in late adolescence (Fig. 4C_2_; Table 1Y). Neither weight differences between the experimental and control animals nor estrous cycle where applicable in females (P38 and 54) correlated with behavior (Supp. Fig. 3). These data demonstrate a remarkable behavioral correlate, specifically social play behavior, to NAc D1r expression patterns in males during adolescent development. While behavior did change over adolescence in females and warrants further investigation, there was no overt correlation with the pattern of D1r expression in the NAc.

**Fig. 4.**
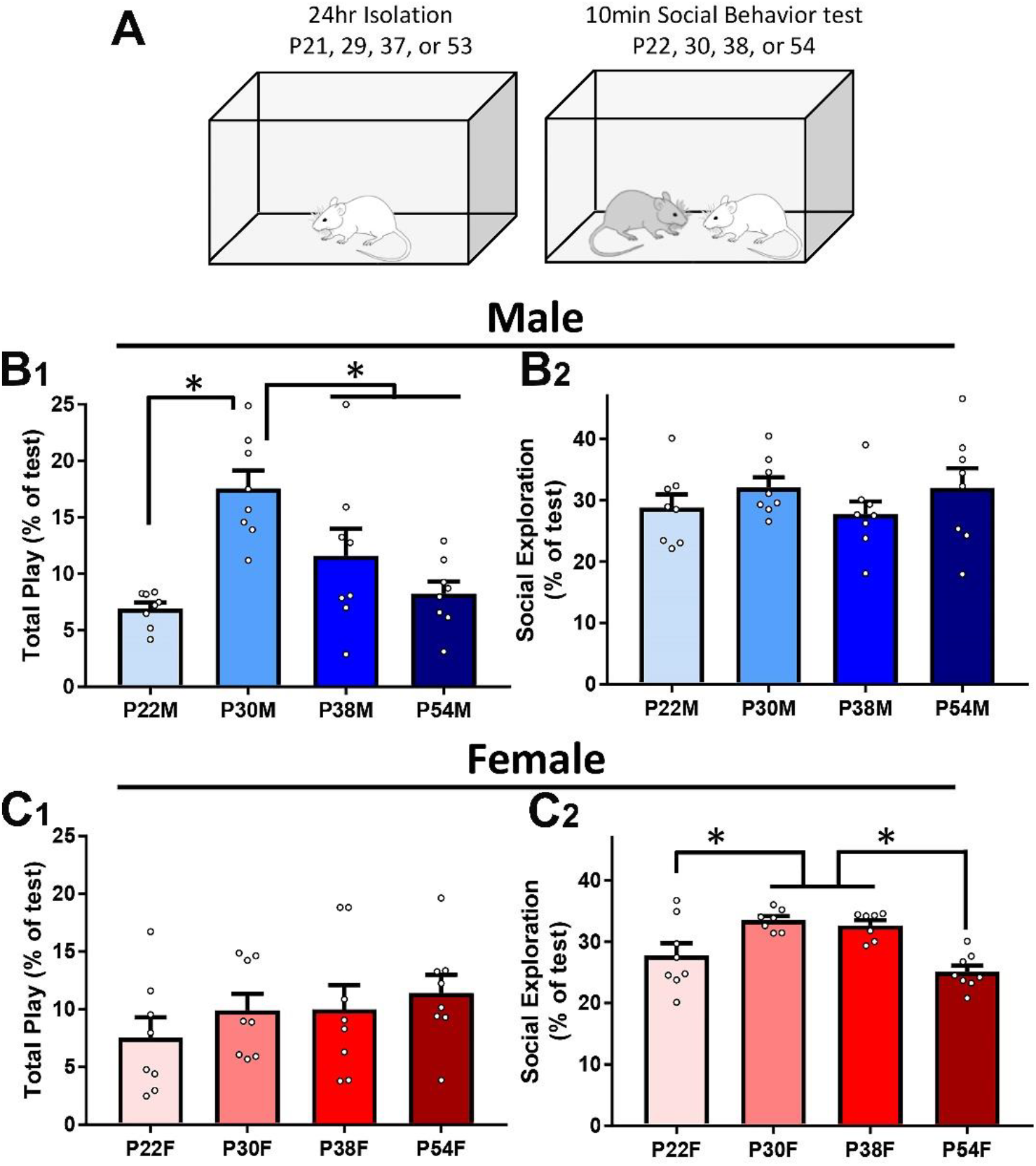
Social behavior patterns in adolescent development are sex-specific, and social play behavior mirrors D1r expression in males, but not females. **(A)** Experimental design: experimental animals (*n*=8/sex/age) were single-housed for 24 hours prior to test, after which time a novel age- and sex-matched conspecific was introduced into their home cage. Ten minutes of interactions were recorded, and later an experimenter blinded to the conditions coded the interactions for either total play or social exploration (i.e. non-play social behavior). Only behaviors initiated by the experimental animal were considered. Data were analyzed with one-way ANOVAs and Holm-Sidak’s post-hoc comparisons. Histograms portray the mean +/− SEM with individual data points overlaid. Significant (*p*<0.05) comparisons are delineated with an asterisk. All statistics are located in Table 1. Males: **(B_1_)** Male social play behavior transiently increased at P30 (Table 1V), **(B_2_)** with no changes in social exploration over development (Table 1W). Females: **(C_1_)** Female social play behavior did not change over development (Table 1X), **(C_2_)** while social exploration peaked from P30-38 (Table 1Y).

In a final set of experiments, we tested the hypothesis that in males, microglial C3R – C3 interactions, demonstrated to be required for D1r elimination in adolescence, are also required for the natural decline that we observed in social play behavior. We bilaterally microinjected 60ng NIF (in 300nL) or vehicle (300nL sterile PBS) into the nucleus accumbens of males at P30, and then assessed their social behavior at P38 (Fig. 5A; Table 1Z). Consistent with our hypothesis, social play behavior (Fig. 5B_1_), but not social exploration (Fig. 5B_2_; Table 1AA), was significantly increased in NIF treated relative to vehicle treated males (also see Supp. Fig. 4). Thus, this report demonstrates for the first time a role for immune signaling and microglia in natural developmental changes in behavior.

**Fig. 5.**
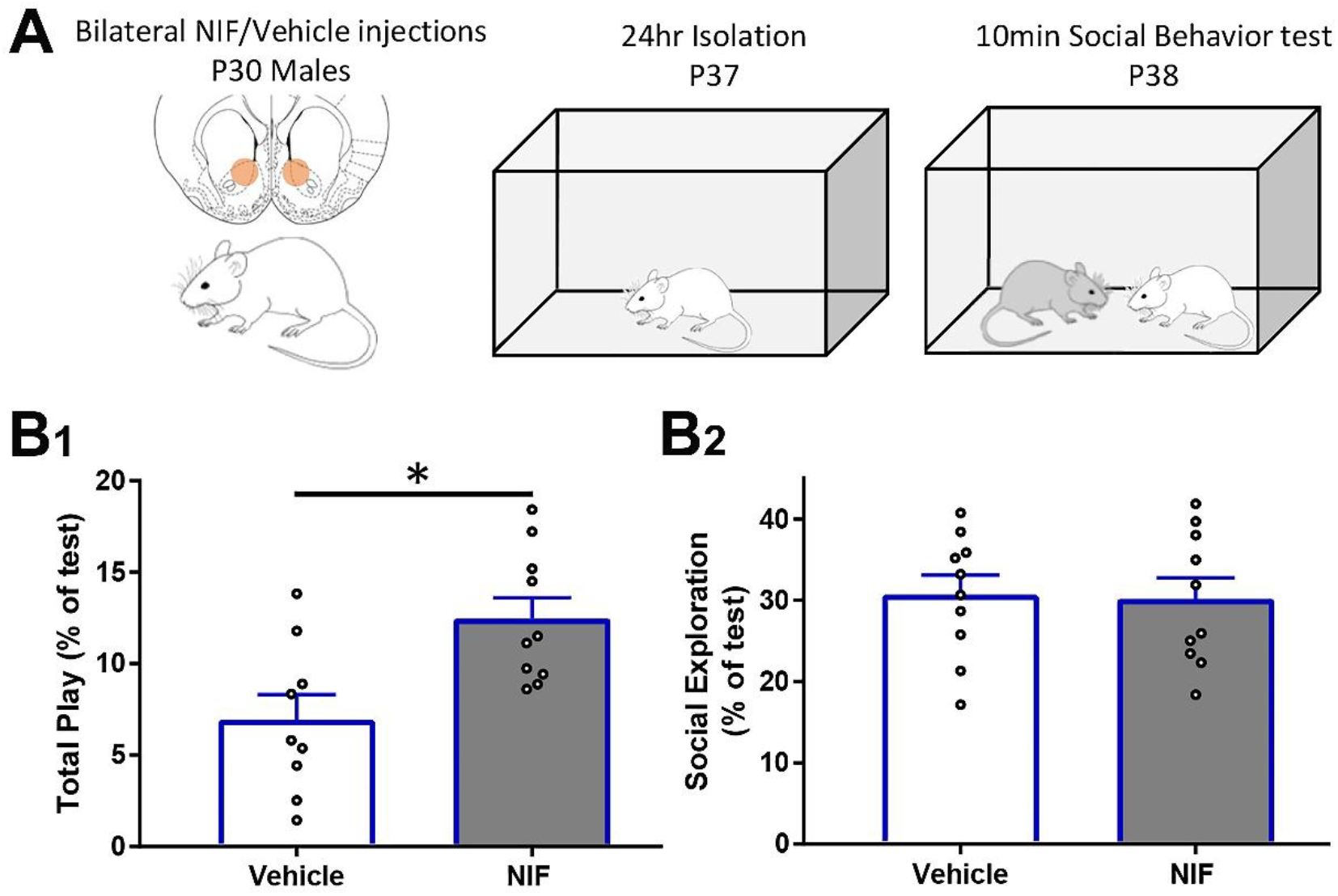
Inhibiting microglial phagocytosis with NIF at P30 in males augments social play behavior later in adolescence. **(A)** Experimental design: P30 males were bilaterally microinjected into the NAc with either NIF or vehicle (*n*=10/treatment). At P37, animals were single-housed for 24 hours prior to test, and then a novel age- and sex-matched conspecific was introduced into their home cage. Ten minutes of interactions were recorded, and later an experimenter blinded to the conditions coded the interactions for either total play or social exploration (i.e. non-play social behavior). Only behaviors initiated by the experimental animal were considered. Data were analyzed with two-tailed unpaired *t*-tests. Histograms portray the mean +/− SEM with individual data points overlaid. Significant (*p*<0.05) comparisons are delineated with an asterisk. All statistics are located in Table 1. **(B_1_)** Male social play behavior at P38 was increased in NIF-treated animals relative to vehicle-treated animals (Table 1Z), **(B_2_)** with no changes in social exploration between treatment groups (Table 1AA).

## Discussion

Herein, we describe a role for microglia and immune signaling in sex-specific NAc development and social behavior during adolescence. D1rs in the NAc are downregulated during adolescent development in males, between early-mid adolescence (P30-38), and in females between the peri-early adolescent period (P20-30). While both sexes undergo D1r downregulation, the mechanisms by which this is accomplished are starkly different: in males, D1rs associated with complement C3 are recognized by microglia via C3R/CD11b, phagocytosed, and lysosomally degraded. However, D1r downregulation in females appears to be independent of complement C3 and microglia. To our surprise, D1r expression patterns in the NAc mirror social play behavior in males (but not females), suggesting that D1r expression (and the immune mechanisms which regulate it) is causally linked with developmental changes in behavior. Indeed, pharmacological interference of microglial C3R (specifically the CD11b subunit) to recognize its ligands, like C3, decreases phagocytic activity *ex vivo*, and increases D1r levels and social play *in vivo*.

The dopaminergic mesolimbic reward circuitry plays a central role in a broad repertoire of cognitive, social, and motor functions, thus linking abnormalities in this system to a wide variety of neurodevelopmental, neuropsychiatric, and neurodegenerative diseases. Intriguingly, a number of neuropsychiatric disorders commonly emerge in adolescence, including depression, anxiety, schizophrenia, and addictive behaviors, many of which present with a striking sex bias.^44,45^ This convergence suggests that there may be instances in which development is misdirected from a normal, healthy state to a disordered, pathologic state, potentially in sex-specific ways. However, the molecular mechanisms underlying reward circuitry development in adolescence are not well known, and, consequently, what causes ‘misdirected development’ or how it is mediated at the molecular level remains unclear.

The neural circuitry underlying social behavior has garnered increasing attention with the recognition of social deficits as a common symptom in a variety of neurodevelopmental (i.e. Autism) and neuropsychiatric (i.e. anxiety or schizophrenia) disorders. Consistent with our data, social play behavior has been reported to be most robust between P30-40 in rats, and generally (though not exclusively) rough-and-tumble play is higher in male than in female adolescents.^46,47^ Interestingly, we did not see changes in female social play behavior; we did, however, observe modest, but significant changes, with little variability, in other social behaviors over adolescence, specifically social exploration, which we defined as allogrooming (social grooming) and conspecific sniffing. Social interactions in adolescence, and play behavior in particular, are thought to be a way for males to ‘practice’ behaviors that result in hierarchical and sexual competency later in life.^48,49^ In females, however, adolescent social interactions may serve roles that we are not adequately assessing. For instance, maternal behaviors and pup interaction is of critical importance to successful adult female behavior in rats, and increased pup-centered behaviors often coincides with adolescence in females more than in males.^50^ Moreover, both the familiarity of a conspecific animal (i.e. a prior cage mate or novel female) as well as the social context (i.e. home cage or novel cage) have been demonstrated to influence female play behavior.^51,52^ Thus, we propose that to understand the molecular mechanisms governing female NAc (and greater reward circuitry) development, we first need to identify the repertoire of experiences that are most *salient to that sex*. The importance of changes in social exploration observed here will provide an excellent foundation on which to begin this exploration.

Many sex differences in neurodevelopment and behavior, including sex-specific social behavior, are established by a surge in male sex hormones (androgens) in male, but not female fetuses around the time of birth.^46,53^ Remarkably, in regions of the brain known to develop highly sexually dimorphic neural architecture, the effects of androgen masculinization at birth is mediated indirectly through microglia and immune signaling.^54,55^ Consistent with the engagement of microglia and immune signaling specifically in the male brain early in life to mediate sexual organization, a recent report in BioRXiv indicates that the human transcriptome in neocortex displays higher phagocytosis- and immune-related gene expression in males than in females prior to birth, which then switches to a higher female bias after birth.^56^ Intriguingly, in this same dataset, there was another age at which males again demonstrated higher immune gene expression relative to females: peri- to early-adolescence. Here, we demonstrate that the male, but not female NAc, engages classical immune signaling to refine dopaminergic circuitry during adolescent development. Collectively, these data raise the possibility that the preferential engagement of immune-mediated mechanisms is pre-programmed, potentially via early-life masculinization events, to serve male development. Consistent with this notion, the mechanisms of male NAc development in adolescence described in this report mirror the role of microglia in synaptic pruning in the male mouse retinogeniculate system,^26,27^ suggesting a ubiquitous role for complement- and microglia-dependent pruning in the developing male brain. Interestingly, several models of neurological disorders including virus-induced cognitive dysfunction,^57^ healthy aging,^58^ and Alzheimer’s disease.^59,60^ Thus, this natural developmental mechanism may be over-activated in pathology, and potentially in sex-specific ways. How this occurs, and what mechanisms might initiate this latent immune process much later in life during adolescence in males, will be important areas to explore. While D1rs are downregulated in the female NAc, there does not appear to be any involvement of the microglial-complement system in this process. Thus female neural development may accomplish largely the same phenotypical and functional outcomes (i.e. appropriate circuit development and behavior), by employing starkly different mechanisms, a phenomena also observed in sex differences in pain processing^61^ and analgesia.^62^

Complement signaling involves a complex cascade of protein activation, either via the “classical,” “lectin,” or “alternative” pathways, all ultimately converging on the cleavage of complement C3.^28,29^ While it is clear that C3 and C3R are critical for neurodevelopment as well as engaged in pathology, how C3 expression and its proteolytic processing is regulated in the brain is unknown. Our data demonstrate that iC3b, identified via Western blotting by a fragment that exhibits a unique molecular weight, can physically associate with NAc D1rs in males and females. Moreover, inhibiting the ability of C3R/CD11b to associate with its ligands, including iC3b, blocks D1r downregulation and augments social play behavior in adolescent males. Importantly, the complement cascade is not the only means of eliminating synapses and refining neural circuitry. Other immune protein families like the major histocompatibility component (MHC) family,^63^ other cell types (astrocytes in particular),^64^ and immune-independent mechanisms (e.g. retraction due to competition^65^ or GABA^A^ receptor-mediated elimination^66^) have all been implicated in synaptic elimination. Interestingly, there is also evidence for neuro-glial communication guiding immune-mediated neural circuit refinement,^67^ and thus the molecular network which regulates C3-D1r mediated elimination is likely to be far more complex than we currently appreciate.

Two important, unexplored lines of inquiry arise from these data. First D1rs are not the only receptor, or even the only dopamine receptor subtype, that is regulated in adolescence. Thus, if C3 and microglia are coordinating the elimination of D1rs in adolescent males, are there other receptors regulated similarly? In particular, D1rs and D2rs, are largely thought to be complementary. While both are G-protein coupled receptors that respond to dopamine, D1rs stimulate cAMP signaling, and thus have an “excitatory” effect in the cell; conversely, D2rs inhibit cAMP production, resulting in an “inhibitory” effect.^68,69^ Thus the ratio of these receptors is important for excitatory:inhibitory balance, and having a complete understanding of the sex-specific regulation of these complementary receptors will be important. And second, there is a well-defined anatomical and functional dissociation between the shell and core of the NAc.^70^ We defined our regions of interest not by core vs. shell, but by D1r immunoreactivity. Thus, while we are likely studying D1r dynamics in the core (i.e. immediately dorso-medial to the anterior commissure), we cannot exclusively claim as much. Interestingly, while core vs. shell distinctions are unquestionably important for some behaviors, dopamine signaling supporting social play did not appear to have a core vs. shell dependency.^17^ Moreover, we focused our analyses primarily to the anterior NAc. There is an interesting anterior-posterior NAc dichotomy, in which the anterior NAc processes reward and posterior NAc processes aversion in humans^71^ and rodents.^72^ Given its different functions and the notion that adolescents experience both reward augmentation and dampened aversion,^73^ it will be important to determine if the development of the NAc is not unitary, but rather utilizing different mechanisms and/or different timescales along the anterior-posterior axis.

Our data raise many important questions, and point to several unexplored lines of inquiry for future exploration. Intriguingly, a recent report demonstrated different microglial signatures within different regions of the reward circuitry,^74^ and understanding how these structures develop in concert and cooperate to mediate behavioral outcomes will be revolutionary for our understanding of adolescent vulnerabilities to neuropsychiatric disorders. While challenging in practice, moving toward an experimental framework that assesses more than one signaling pathway and/or brain regions in parallel can reveal unappreciated cooperation in the service of behavior,^75^ and will ultimately be required to fully understand the development of this system. We thus view our data as the tip of the iceberg, and provide a foundation on which to initiate in-depth molecular dissection of the impact of adolescent experience on reward circuitry development and dysfunction that we propose will be informative for many different aspects of neuroscience.

## Acknowledgements

We thank Phuong K. Tran and Ernesto Barbosa for help with experiments, and members of the Bilbo lab for helpful discussion during the development of this project. This work was supported by RO1 DA034185 and RO1 MH101183 to SDB and F32DA043308 to AMK.

## Author Contributions

AMK, CJS, and SDB planned the experiments. AMK, CJS, NRA, and SCS performed experiments. AMK analyzed the data and wrote the manuscript. All authors contributed to manuscript edits.

## Methods

### Animals

Adult male and female Sprague-Dawley rats (both age <u>P</u>ostnatal day 70-75 (P70-75); Harlan/Envigo, Dublin, VA) were purchased to be breeding pairs and pair-housed in individually ventilated cages with *ad libitum* access to food (Purina lab Diet 5001) and water. The colony was maintained at 23°C on a 12:12 light-dark cycle (lights on at 0700 hours) and cages were changed once per week. Females were separated from males after 20 days for birth. Litters were culled to a maximum of 12 pups on P2-5, and at P21 pups were weaned into same sex pair-housing (2-4 rats per cage). All experiments were approved by the Massachusetts General Hospital Institutional Animal Care and Use Committee.

### Tissue Collection

Animals were euthanized at the ages indicated in each experiment by CO2 anesthesia and exsanguination. Blood was collected prior to exsanguination via cardiac puncture, and then 0.9% saline was perfused until blood was cleared. Animals were decapitated and brains extracted. For co-immunoprecipitation experiments, bilateral nucleus accumbens was dissected, frozen in isopentane, and stored at −80C until processing. For immunohistochemical experiments, brains were post-fixed in 4% paraformaldehyde (PFA) in PBS. In a subset of surgically manipulated male and all surgically manipulated female subjects, euthanasia was performed as previously described, with 4% PFA following saline perfusion. Whole brains were then incubated in 4% PFA for 24 hrs at 4°C. The remaining surgically manipulated male subjects were subjected to social play testing, and then euthanized as above 90 mins after behavioral testing.

### Immunohistochemistry

PFA-fixed brains were cryoprotected in 30% sucrose in 0.1M PB for at least 2 days. Cryoprotected brains were rapidly frozen in molds with Tissue Tek on dry ice, and then cryosectioned at 20μm. Four animals from different litters were represented in each experimental group. Sections were stored in 0.1M PB with sodium azide (0.1%) until processing. After washing in PBS, sections were incubated at 80°C for 30 mins in 10mM citric acid (pH 9.0) for epitope retrieval. Background fluorescence was quenched sequentially with 30 min incubations in 1mg/mL sodium tetraborate in 0.1M PB and then 50% methanol. Tissue was permeabilized for 1 hr with 0.3% Triton-x100 in PBS and then blocked in 10% normal donkey serum (Jackson ImmunoResearch; West Grove, PA) in PBS for 1 hr. Primary antibodies were applied sequentially for overnight incubations in 5% normal donkey serum in PBS at 4°C: either anti-goat C3 (MP Biomedicals #0855713; Santa Ana, CA; 1:200), anti-mouse D1r (Novus Biologicals #NB110-60017; Littleton, CO; 1:750), anti-rabbit Iba1 (Wako Chemicals #019-19741; Richmond, VA; 1:1500) or anti-rabbit CD68 (Abcam #ab125212; Cambridge, MA; 1:5000), anti-goat C3, anti-mouse D1r. Secondary antibodies were applied simultaneously for the C3-D1r-Iba1 triple stain (Donkey anti-goat Alexa Fluor (AF) 647, anti-mouse AF 488, anti-rabbit AF 563; Thermo Fisher Scientific, each 1:500) for 2hrs at room temperature in the dark. For the CD68-C3-D1r triple stain, CD68 was amplified prior to proceeding with the subsequent primary incubations. Tissue was incubated with biotinylated anti-rabbit (Vector Laboratories; Burlingame, CA; 1:1000) for 2hrs at room temperature, in Avidin-Biotin Complex solution for 30min at room temperature (Vector Laboratories; Burlingame, CA), and then Tyramide Signal Amplification fluorescein reagent (PerkinElmer) for 10mins at room temperature in the dark. After washing, the tissue was then incubated sequentially with C3 and D1r antibodies, and then incubated with secondary antibodies (anti-goat AF 563, anti-mouse AF 647; Thermo Fisher Scientific, 1:500) for 2hrs at room temperature in the dark. All tissue was mounted on gelatin subbed slides, coverslipped with ProLong Antifade Mounting media (Thermo Fisher Scientific), sealed with nailpolish, and stored at −20°C until imaging.

### Image Acquisition and Volumetric Reconstructions

One to three z-stacks (depending on the size of the ROI, see Supp. Fig. 1A) per section within the D1r-rich region of the nucleus accumbens were acquired on a Nikon A1SiR confocal microscope with 60x magnification and 0.1μm step size (*n*=4 animals per group, 2 sections per animal). Male and female tissue required different imaging parameters and thus are analyzed and graphically depicted separately. While exhibiting the same visual features (i.e. rhinal fissure size and corpus callosum shape) used to select the NAc, female ROIs tended to be smaller than males, and thus fewer z-stacks were taken from female tissue (specific differences in Table 1). Background was subtracted and threshold values were recorded using ImageJ (National Institutes of Health; Bethesda, MD). These files were then uploaded into Imaris (Bitplane; Zurich, Switzerland) to create volumetric reconstructions of (i) each individual channel, (ii) C3-D1r colocalization, and (iii) ‘masked’ C3, D1r, and C3-D1r. The masking application refers to the use of one channel as a 3D outine, or mask, on which all other channels are superimposed. If other channels all within this 3D mask, they are considered to be in contact with or contained within the masking channel. For example, Iba1-masked D1r volume would constitute all D1r volumes that are either within (i.e. engulfed) or directly contacted by microglia, whereas a CD68-masked D1r volume constitutes all D1r volumes within a CD68+ lysosome. Total volumes for each channel reconstructed in Imaris (a maximum of 24 values across 4 different animals per group) were then used for statistical analysis.

### Co-Immunoprecipitation (Co-IP)

Bilateral NAc were dissected from one animal/sex/age group. Pilot experiments yielded very low D1r reactivity in homogenate, so we performed crude membrane fractionation prior to Co-IP. Tissue was homogenized with a Tissue Tearor at the lowest setting (~5000 r.p.m.; Biospeck Products; Bartlesville, OK) in 1mL 0.32M sucrose, 1mM HEPES, 50mM NaF, 1mM activated NaOv, and protease inhibitors (cOmplete Mini, EDTA-free #1183617001; Roche Diagnostics; Mannheim, Germany). Homogenate was centrifuged at 1000xg for 10 mins at 4°C to pellet debris and nuclei, and the supernatant was centrifuged at 12,000xg for 20 mins at 4°C (these parameters are the same for future references of centrifugation). The pellet was reconstituted in sucrose buffer, and 5% was removed for input BCA assessment. All samples were centrifuged, and pellets were reconstituted in 1% PFA in sucrose buffer and incubated at room temperature for 10 mins. Glycine in sucrose buffer (250mM) was added to quench the crosslinking reaction and samples were centrifuged. Pellets were resuspended in Co-IP buffer (20mM Tris pH 7.4, 50mM NaCl, 2mM EDTA, 1% CHAPS, 1mM activated NaOv, 50mM NaF with protease inhibitors) and homogenized gently until fully solubilized. After normalizing protein levels (based off of input BCA assay assessment; Thermo Fisher Scientific), 3µg of anti-mouse D1r antibody (Novus Biologicals #NB110-60017; Littleton, CO; 1:750) or control mouse IgG (Santa Cruz Biotechnology #sc-3879; Dallas, TX) were incubated with samples overnight. Protein G DynaBeads (Thermo Fisher Scientific) were washed with and then reconstituted in Co-IP buffer, and then samples were incubated with 25µL DynaBeads at 4°C for 2 hrs. Samples were placed on magnets, and beads were washed with Co-IP buffer. Beads were then incubated in high-salt lysis buffer (200mM NaCl, 100mM Tris (pH 8.0), 20mM EDTA, 50mM NaF, 1mM activated NaOv, 5% SDS with protease inhibitors) and incubated at 65°C for 4 hrs to reverse crosslinking and elute antibody-protein complexed from the DynaBeads. Sample buffer was added, and samples were boiled for 10mins. Western blots were performed as in Kopec et al.^76^ Briefly, samples were loaded onto NuPAGE 4-12% 1.0 mm Bis-XX gels (Thermo Fisher Scientific) for electrophoretic separation, transferred to 0.22µm nitrocellulose membranes (LI-COR Biosciences; Lincoln, NE), blocked with LI-COR PBS blocking buffer (LI-COR Biosciences; Lincoln, NE), and incubated with primary antibodies (1:500 anti-rabbit D1r, EMD Millipore #AB1765P; 1:500 anti-rabbit PSD95, Thermo Fisher Scientific #51-6900; anti-rabbit gephyrin, Abcam #ab32206; 1:250 anti-goat C3, MP Biomedicals #0855713) overnight at 4°C. After washes, secondary antibodies (LI-COR infra-red goat anti-rabbit or donkey anti-goat 800CW, LI-COR Biosciences; Lincoln, NE) were incubated for 1hr at room temperature, and bands were visualized using an Odyssey LI-COR imaging system (LI-COR Biosciences; Lincoln, NE).

### In Vitro Phagocytosis Assay

Microglia were isolated from minced female brains (n=4) according to XX. Briefly, tissue was enzymatically and mechanically disrupted and then myelin and debris removed via Percoll gradient centrifugation. Resulting cell bodies were incubated with anti-rat CD11b (i.e. microglia-specific) magnetic beads (Miltenyi Biotec #130-105-634; Auburn, CA) and then separated from the CD11b-population via magnetic columns. CD11b+ cell numbers were estimated on a hemocytometer, and 75,000 cells per well were incubated on 12mm glass coverslips (Thermo Fisher Scientific #12-545-80) in media (1% L-glutamine, 1% Pen-Strep, 1% N2 media, 1% sodium pyruvate, 0.1% forskolin in DMEM) at 37C/XX CO2 for 90 minutes with either vehicle, 60ng (1x reconstitution; 200μg/mL in PBS) neutrophil inhibitory factor (NIF), or 120ng (2x reconstitution; 400μg/mL in PBS) NIF (R&D Systems #5845-NF-050; Minneapolis, MN). Each condition was performed in duplicate. Fluorescbrite YG carboxylate microspheres (1μm diameter; Polysciences, Inc.; Warrington, PA) were added 1:1000 to each well and plates were re-incubated for 90 mins. Immunocytochemistry was performed as in Derecki et al.^77^ Coverslips were fixed with 4% PFA for 20 mins at room temperature, and then washed in PBS. Coverslips were blocked and permeabilized for 1 hr at room temperature (10% normal donkey serum, 0.5% BSA, 0.3% Triton x-100 in PBS), and then incubated with rabbit anti-Iba1 antibody (1:700; Wako Chemicals #019-19741; Richmond, VA) overnight at 4C. After washing in PBS, coverslips were incubated with donkey anti-rabbit AF 563 (1:500; Thermo Fisher Scientific) for 2 hrs at room temperature, washed, and mounted with Pro-Long Antifade Mounting media with DAPI (Thermo Fisher Scientific). Images comprised of DAPI, TexasRed (Iba1), and GFP (beads) were acquired at 20x magnification with a Zeiss AxioImager microscope equipped with a z-drive, Apotome optical dissector, and AxioCam HRm (Carl Zeiss Inc., Gottingen, Germany). Three 3-step (1μm) z-stacks were acquired for each coverslip, and average z-projections were created using ImageJ (National Institutes of Health; Bethesda, MD). Cell bodies were outlined by an experimenter blinded to the conditions using a merged DAPI and Iba1 image in the ROI manager of ImageJ. Densitometric measurements of the GFP channel (beads), were then acquired using the pre-set ROIs. One value (the average integrated GFP density for cells in each image) for each of 3 z-stacks, for each of 2 coverslips, for each of 4 animals, resulted in 24 data points.

### Stereotaxic Microinjection Procedure

Male (P30) and female (P22) rats were maintained under isofluorane anesthesia for the entire surgical procedure (2-3%; VetEquip; Livermore, CA). The scalp was cut midsagittally and Bregma was marked, after which two bilateral holes drilled at AP +2.25mm, ML ±2.5mm, DV −5.75mm coordinates in males, or AP +2.7mm, ML ±2.4mm, DV −5.55mm coordinates in females. A Hamilton syringe (Hamilton #7105; Reno, NV) was lowered to depth at a 10° angle and left in place for 1 min. NIF (1x reconstitution, 200μg/mL NIF; R&D Systems; Minneapolis, MN) or vehicle (sterile PBS) was injected at a rate of 50nL/min (60ng in 300nL in males, 50ng in 250nL in females). The syringe was left in place for 5min for diffusion, and then retracted and the procedure repeated on the other hemisphere (see Supp. Fig. 2). Saline was used to wet the scalp, and then the wound was closed with surgical staples and coated topically with Bupivicaine. An injection of Ketofen (5mg/kg) was administered subcutaneously and the animal was placed in a clean cage with a food pellet and gelatinized water for recovery. After 2-3 hrs, the animal was re-paired with its cagemate and monitored for the next 48 hours. Right versus left hemisphere and drug versus vehicle injections were counterbalanced, and the syringe was thoroughly cleaned with cleaning solution (Hamilton cleaning concentrate, Hamilton; Reno, NV) and distilled water before being used for the next surgery, or in the case of within-animal vehicle vs. NIF treatment, between injections. Injection coordinates and volumes were estimated with bromophenol blue dye in water and immediate decapitation.

### Social Behavior Assessment

Animals were handled on 3 occasions (~5min per session) prior to any behavioral tests. Experimental animals were single-house for 24 hours at age P21, P29, P37, or P53, while age- and sex-matched conspecifics were always group-housed (2-3 animals/cage). Social play tests were performed in the three hours prior to the dark cycle (1600-1900) as follows: experimental and conspecific animals were each weighed and then the conspecific animal was placed in the experimental animal’s cage. Ten minutes of behavior was recorded using ANY-maze software (ANY-maze Behavioural tracking software; Wood Dale, IL), after which the conspecific animal was returned to its cage. When testing females, the experimental animal was inspected for a vaginal opening, and if appropriate, estrous smears were taken and later classified as metestrus, diestrus, proestrus, estrus phases of the estrous cycle according to Goldman et al.^78^ Behavior initiated by the experimental animal over the first 9 minutes of the test was scored by a blinded investigator using Solomon Coder software (András Péter; solomoncoder.com). Social behaviors were determined as described in Manduca et al.^17^:

i. Percentage of time spent in social play, which included any classical play posturing (i.e. nape attacks, supine, and pins) in addition to pouncing on the conspecific, play boxing, and any rough-and-tumble play bouts.
ii. Percentage of time spent in social exploration, which included grooming or sniffing the conspecific animal.

Conspecific animals were never litter mates with and no more than +/−2 days different in age from the experimental animal. Weight differences between the animals varied, with surgical intervention generally causing increased body weight relative to non-surgically manipulated animals at the same age. However, difference in body weight did not correlate with total play scores (Supp. Fig. 3,4) in males or females at any age or experimental group.

### Statistical Analysis

#### Immunohistochemistry

Each target protein (D1r, C3, and C3-D1r) was calculated as a percent of the total volume (in the case of different z-stack sizes). To assess microglial contact, masked volumes were calculated as a percentage of the total possible volume (i.e. masked D1r divided by total D1r), and then divided by the percentage of Iba1 volume in the z-stack to account for changing levels of protein. To assess association with CD68+ lysosomes, masked protein was calculated as a percentage of total possible volume. One-way ANOVAs were performed for all developmental time course data (i.e. P20, 30, 38, 54). In the event of significant ANOVAs, post-hoc Holm-Sidak *t*-tests were conducted (two-tailed). Because (i) immunofluorescent data needed to be acquired at different imaging parameters for males and females and (ii) we were interested sex differences in patterns of expression and behavior over development, and not sex differences at any one age, we did not incorporate sex as an independent variable, and rather performed a one-way ANOVAs for each sex. For the analysis of D1r expression in NIF- and vehicle-treated NAc within the same animals, paired *t*-tests (two-tailed) were conducted.

#### Behavior

Duration of social play and social exploration were calculated as a percent of the total testing period (500-540s), and then one-way ANOVAs were conducted separately for males and females. When appropriate, Holm-Sidak post-hoc *t*-tests (two tailed) were conducted. In bilateral NIF- and vehicle-treated males, behavior was analyzed with unpaired *t*-tests (two-tailed).

Data points greater than two standard deviations above the group mean were excluded as outliers. All data were analyzed, and figures created with, GraphPad Prism 7 (GraphPad Software; La Jolla, CA). Data are depicted in histograms as average ± standard error of the mean with individual data points overlaid. All statistics are located in Table 1.

## Supplementary Materials

**Supp. Fig. 1.**
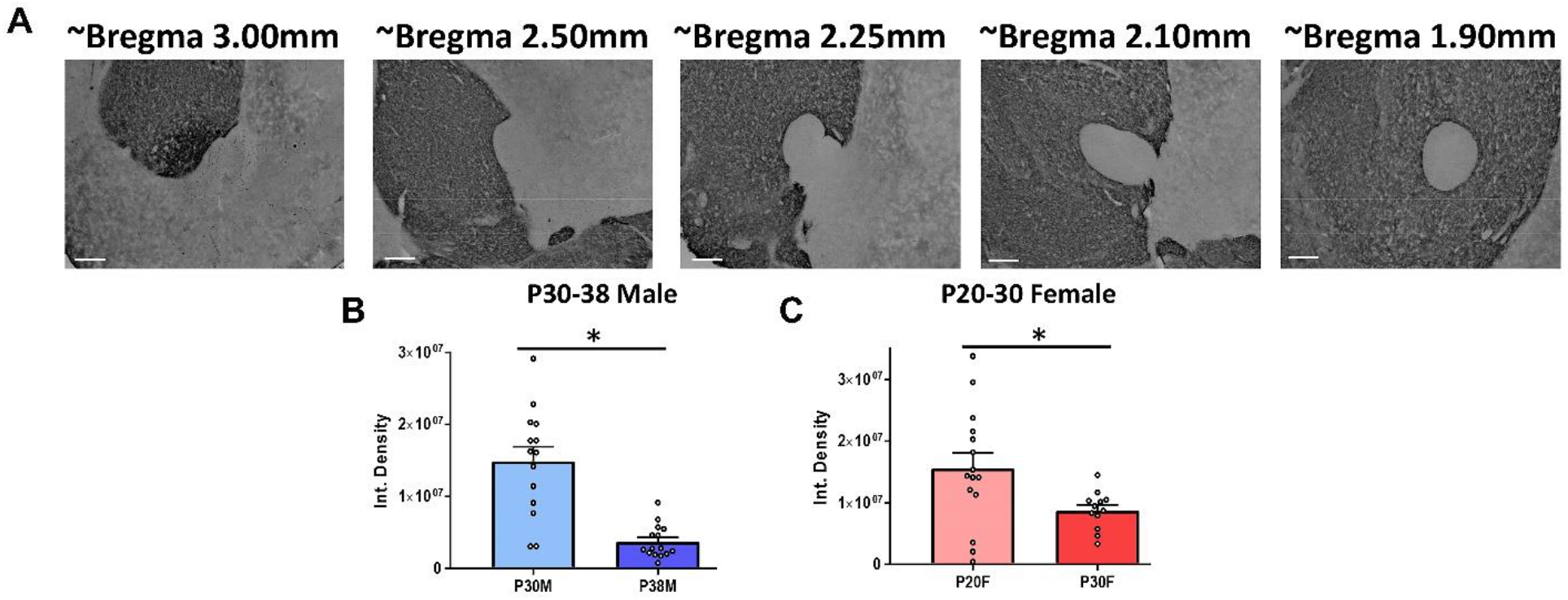
D1r immunoreactivity in the dorsomedial NAc. **(A)** Chromogenic immunohistochemistry of D1r reactivity in the NAc (example images are male). In all images, the midline is toward the left and dorsal brain is toward the top. In immunofluorescent analyses (Fig. 1-4), z-stacks were acquired in the dorso-medial D1r enriched areas of the NAc. **(B)** D1r downregulation between P30-38 in males (*t*_(27)_=5.561, *p*<0.0001) **(C)** P20-30 in females (*t*_(25)_=2.408; *p*=0.024) was confirmed (Fig. 1) in a separate cohort (*n*=3 animals/sex/age). Data were analyzed with two-tailed unpaired *t*-tests. Histograms portray the mean +/− SEM with individual data points overlaid. Significant (*p*<0.05) comparisons are delineated with an asterisk.

**Supp. Fig. 2.**
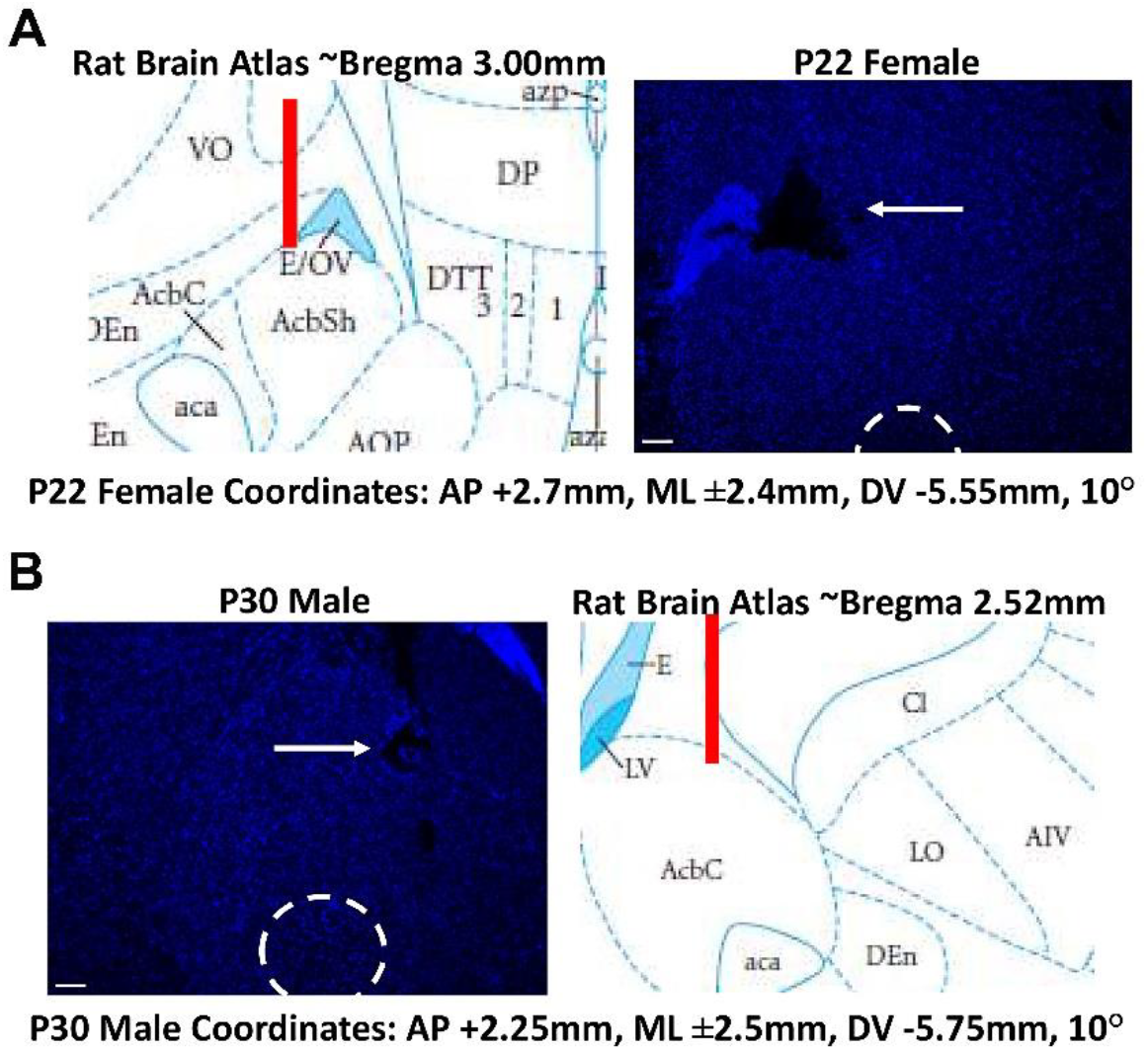
Microinjection coordinates and estimates of stereotaxic manipulation. Stereotaxic coordinates and volumes were estimated by injecting bromophenol blue dye into the brain under high levels of anesthesia (non-survival), immediately sacrificing the animal, and taking coronal sections through the brain. The microinjection technique was selected for these experiments to reduce the level of inflammatory (i.e. microglial) reaction to the manipulation, as we hypothesized there was a an on-going, natural immune process occurring in adolescent development. Brain healing was so effective that microinjection track marks were undetectable in the vast majority of experimental animals (8-10 days after injection). Because we are uncertain that the dye used to estimate coordinates under very high anesthesia would recapitulate dynamics of NIF/vehicle injection under lower anesthesia (survival surgeries), we microinjected NIF bilaterally in males and females, allowed the animals to wake and resume normal behavior, and then sacrificed the animals 2-4 hours later. Example microinjection tracts for a P22 female **(A)** and P30 male **(B)** are visualized with DAPI to help detect neural architecture and comparable locations from a Rat Brain Atlas are aligned. In both sexes, the tract targets the dorso-medial NAc (see Supp. Fig. 1 for comparable D1r immunoreactivity).

**Supp. Fig. 3.**
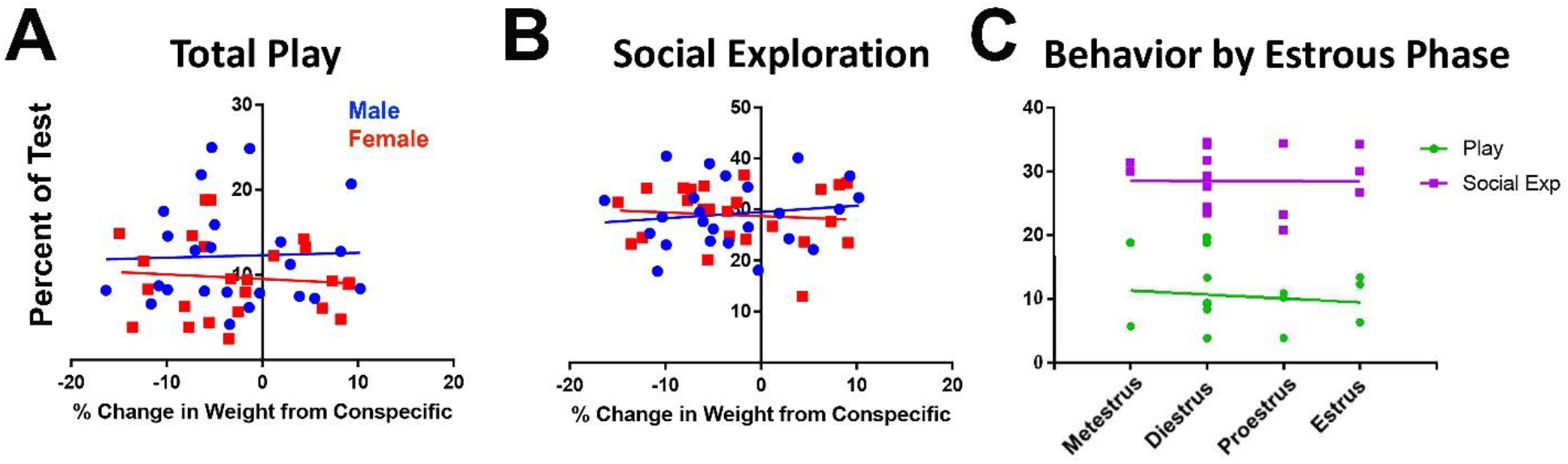
Social behavior does not correlate with weight differences between the experimental and conspecific sex in either males or females, nor estrous phase in P38-54 females. **(A-B)** Change in weight between the experimental and conspecific animals was calculated as a percent change from conspecific (Experimental-Conspecific / Experimental * 100), and then plotted against total play or social exploration scores. **(A)** Pearson correlations between weight and total play were nonsignificant for males (Pearson’s *r*=0.03) and females (Pearson’s *r*=0.69). **(B)** Pearson correlations between weight and social exploration were nonsignificant for males (Pearson’s *r*=0.14) and females (Pearson’s *r*=0.09). **(C)** If experimental females had vaginal openings (only 1 female at P30 and all females at P38 and P54), estrous smears were collected and classified as metestrus, diestrus, proestrus, or estrus via light microscopy according to XXX. Total play and social exploration behavior scores were then plotted against these four categories. Pearson correlations between estrous phase and total play (Pearson’s *r*=-0.11) and social exploration (Pearson’s *r*==0.004) were nonsignificant.

**Supp. Fig. 4.**
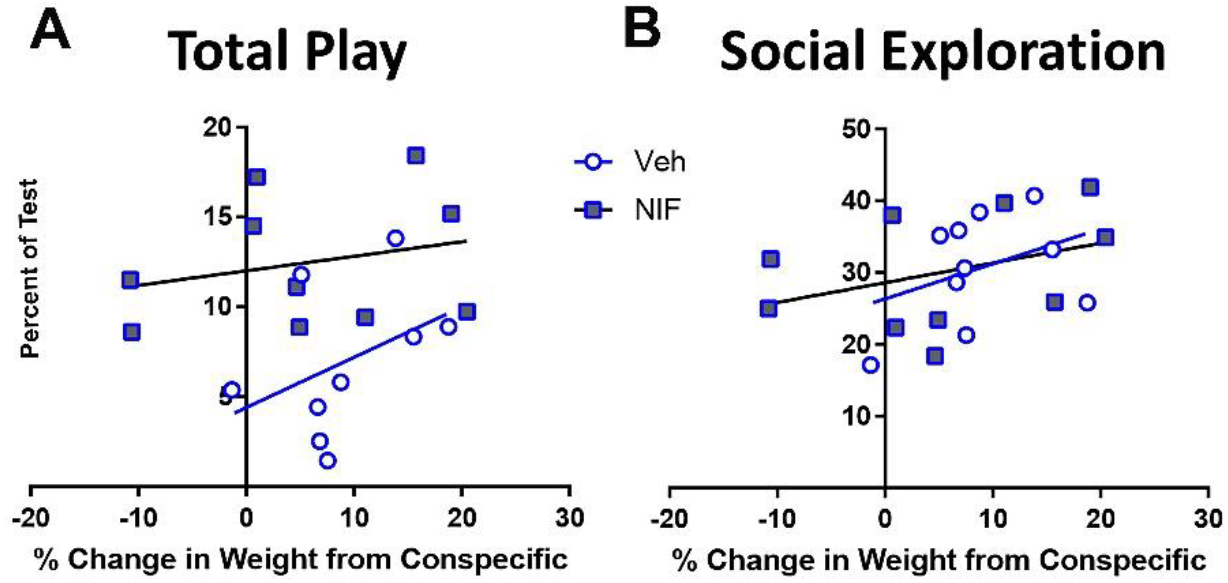
Social behavior assessed in NIF or vehicle microinjected males does not correlate with weight differences between the experimental and conspecific sex. Change in weight between the experimental and conspecific animals was calculated as a percent change from conspecific (Experimental-Conspecific / Experimental * 100), and then plotted against total play or social exploration scores. **(A)** Pearson correlations between weight and total play were nonsignificant for vehicle-injected (Pearson’s *r*=0.41) and NIF-injected (Pearson’s *r*=0.25) males. **(B)** Pearson correlations between weight and social exploration were nonsignificant for vehicle-injected (Pearson’s *r*=0.14) and NIF-injected (Pearson’s *r*=0.14) males.

